# The Spike D614G mutation increases SARS-CoV-2 infection of multiple human cell types

**DOI:** 10.1101/2020.06.14.151357

**Authors:** Zharko Daniloski, Tristan X. Jordan, Juliana K. Ilmain, Xinyi Guo, Gira Bhabha, Benjamin R. tenOever, Neville E. Sanjana

**Affiliations:** New York Genome Center, New York, NY, USA; Department of Biology, New York University, New York, NY, USA; Department of Microbiology, Icahn School of Medicine at Mount Sinai, New York, NY, USA; Department of Cell Biology and Skirball Institute of Biomolecular Medicine, New York University School of Medicine, New York, NY, USA

## Abstract

A novel isolate of the SARS-CoV-2 virus carrying a point mutation in the Spike protein (D614G) has recently emerged and rapidly surpassed others in prevalence. This mutation is in linkage disequilibrium with an ORF1b protein variant (P314L), making it difficult to discern the functional significance of the Spike D614G mutation from population genetics alone. Here, we perform site-directed mutagenesis to introduce the D614G variant and show that in multiple cell lines, including human lung epithelial cells, that the D614G mutation is up to 8-fold more effective at transducing cells than wild-type. We demonstrate increased infection using both Spike-pseudotyped lentivirus and intact SARS-CoV-2 virus. Although there is minimal difference in ACE2 receptor binding between the Spike variants, we show that the G614 variant is more resistant to proteolytic cleavage *in vitro* and in human cells, suggesting a possible mechanism for the increased transduction. This result has important implications for the efficacy of Spike-based vaccines currently under development in protecting against this recent and highly-prevalent SARS-CoV-2 isolate.

---

Recently, a novel isolate of the SARS-CoV-2 virus carrying a point mutation in the Spike protein (D614G) has emerged and rapidly surpassed others in prevalence, including the original SARS-CoV-2 isolate from Wuhan, China. This Spike variant is a defining feature of the most prevalent clade (A2a) of SARS-CoV-2 genomes worldwide (Bhattacharyya et al., 2020; Hadfield et al., 2018). Using phylogenomic data, several groups have proposed that the D614G variant may confer increased transmissibility leading to positive selection (Bhattacharyya et al., 2020; Korber et al., 2020), while others have claimed that currently available evidence does not support positive selection (Dorp et al., 2020). Furthermore, in the A2a clade, this mutation is in linkage disequilibrium with a ORF1b protein variant (P314L) (Bhattacharyya et al., 2020), making it difficult to discern the functional significance of the Spike D614G mutation from population genetics alone.

Here, we perform site-directed mutagenesis on a human codon-optimized spike protein to introduce the D614G variant (Shang et al., 2020) and produce SARS-CoV-2-pseudotyped lentiviral particles (S-Virus) with this variant and with D614 Spike. We show that in multiple cell lines, including human lung epithelial cells, that S-Virus carrying the D614G mutation is up to 8-fold more effective at transducing cells than wild-type S-Virus. Similar experiments using intact SARS-CoV-2 further confirms that Spike G614 leads to higher viral infection of human cells. Although we find minimal differences in ACE2 receptor binding between the Spike variants, we show that the G614 variant is more resistant to cleavage *in vitro* and in human cells, which may suggest a possible mechanism for the increased transduction. Given that several vaccines in development and in clinical trials are based on the initial (D614) Spike sequence (Lurie et al., 2020; Yu et al., 2020), this result has important implications for the efficacy of these vaccines in protecting against this recent and highly-prevalent SARS-CoV-2 isolate. For example, neutralizing antibodies that target the receptor binding domain seem largely unaffected in potency but it remains to be seen whether the D614G variant alters neutralization sensitivity to other classes of anti-Spike antibodies (Yurkovetskiy et al., 2020).

The first sequenced SARS-CoV-2 isolate (GenBank accession MN908947.3) and the majority of viral sequences acquired in January and February 2020 contained an aspartic acid at position 614 of the Spike protein (**Figure 1a**). Beginning in February 2020, an increasing number of SARS-CoV-2 isolates with glycine at position 614 of the Spike protein were identified. We found that ∼72% of 22,103 SARS-CoV-2 genomes that we surveyed from the GISAID public repository in early June 2020 contained the G614 variant (Shu and McCauley, 2017). Previously, Cardozo and colleagues reported a correlation between the prevalence of the G614 variant and the case-fatality rate in individual localities using viral genomes available through early April 2020 (Becerra-Flores and Cardozo, 2020). Using a ∼10-fold larger dataset, we found a smaller yet significant positive correlation between the prevalence of G614 in a country with its case-fatality rate (*r* = 0.29, *p* = 0.04) (**Figure 1b**). There has been little consensus on the potential function of this mutation and whether its spread may or may not be due to a founder effect (Bhattacharyya et al., 2020; Dorp et al., 2020). Recently, two separate groups at the University of Sheffield and at the University of Washington have found that in COVID-19 patients there is a ∼3-fold increase in viral RNA during quantitative PCR-based testing for those patients with the G614 variant (Korber et al., 2020; Wagner et al., 2020) (**Figure 1c, d**). Although there is a consistent difference in qPCR amplification between the sites (∼5 C_t_) potentially due to different sampling procedures, RNA extraction methods, qRT-PCR reagents or threshold cycle settings (**Figure 1c**), the difference in amplification (ΔΔC_t_) between G614 and D614 variants is remarkably consistent (1.6 C_t_ for Sheffield, 1.8 C_t_ for Washington), suggesting that this may be due to a biological difference between COVID-19 patients with specific Spike variants (**Figure 1d**).

**Figure 1.**
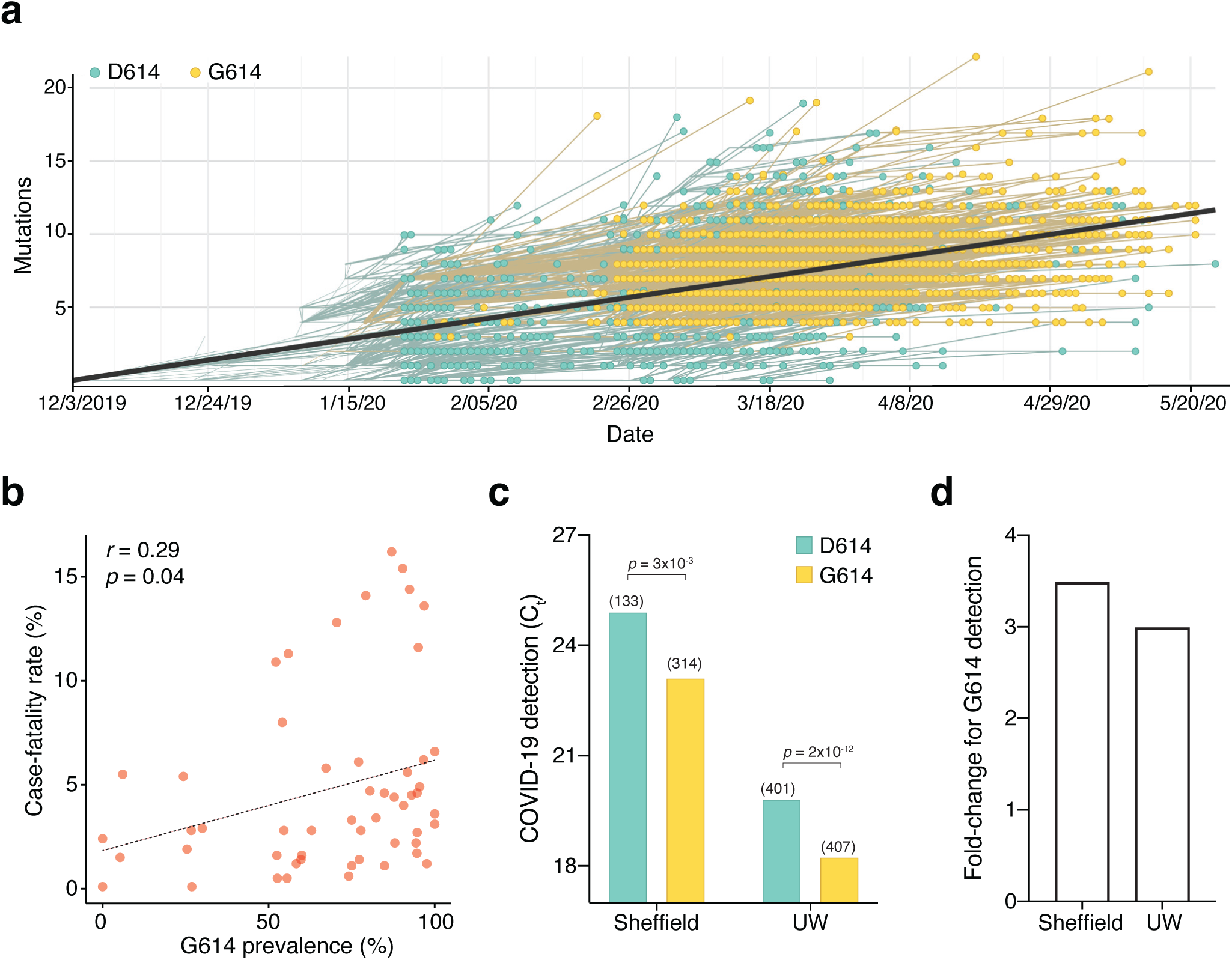
The SARS-CoV-2 D614G mutation has spread rapidly and is correlated with increased fatality and higher viral load. (**a**) Prevalence of D614G-containing SARS-CoV-2 genomes over time. This visualization was produced by the Nextstrain webtool using GISAID genomes (*n* = 3,866 genomes samples from January 2020 to May 2020). (**b**) Per-country correlation of G614 prevalence versus the case-fatality rate (*n* = 56 countries and 22,103 genomes). (**c**) Threshold cycle for quantitative polymerase chain reaction (qPCR) detection of COVID-19 from patients with D614 and G614 Spike. Numbers in parentheses indicate the number of COVID-19 patients in each group and significance testing is using the Wilcoxon rank sum test. This Sheffield data was originally presented in Korber et al. (2020). The University of Washington data was originally presented in Wagner et al. (2020). (**d**) Fold-change of increase in viral RNA present in COVID-19 patient samples with G614 Spike as compared to those with D614 Spike.

Given these findings, we wondered whether the G614 variant may confer some functional difference that impacts viral transmission or disease severity. To address this question, we used a pseudotyped lentiviral system similar to those developed previously for SARS-CoV-1 (Moore et al., 2004). Using site-directed mutagenesis and a human-codon optimized SARS-CoV-2 spike coding sequence (Shang et al., 2020), we constructed EGFP-expressing lentiviruses either lacking an attachment protein or pseudotyped with D614 Spike or G614 Spike (**Figure 2a**). After production and purification of these viral particles, we transduced human cell lines derived from lung, liver and colon. Others have observed increased S-virus transduction in cells that overexpress the angiotensin-converting enzyme 2 (ACE2) receptor (Li et al., 2003; Moore et al., 2004); we also found that S-virus is much more efficient at transducing human cell lines when the human ACE2 receptor is overexpressed (**Supplementary Fig. 1**). Given this, for two of the human cell lines (A549 lung and Huh7.5 liver), we overexpressed the ACE2 receptor to boost viral transduction.

**Figure 2.**
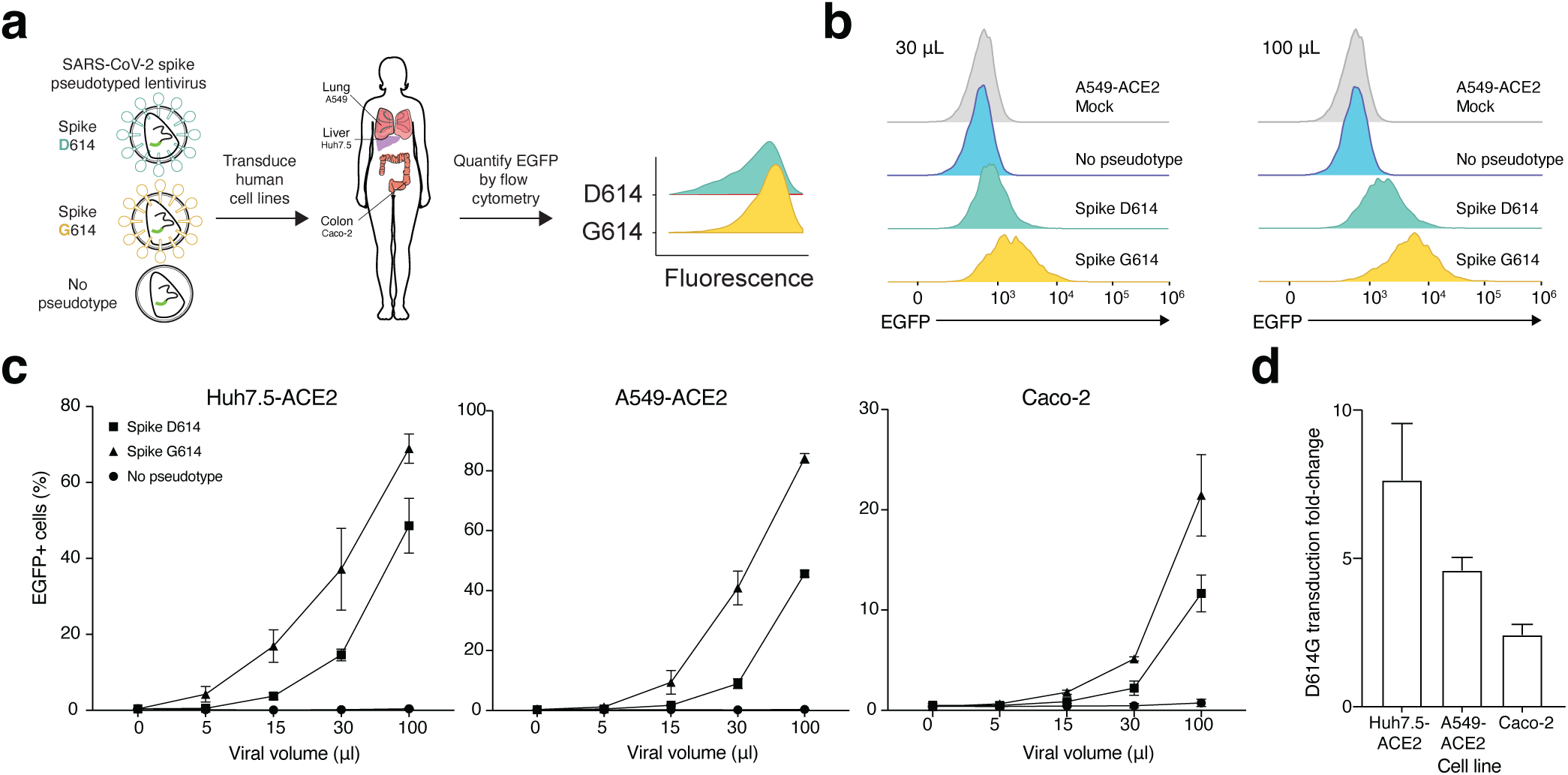
SARS-CoV-2 Spike D614G pseudotyped lentivirus results in increased transduction of human lung, liver and colon cell lines. (**a**) Schematic of EGFP lentivirus pseudotyped with SARS-CoV-2 Spike proteins (or no pseudotype) and readout of EGFP fluorescence by flow cytometry. (**b**) Flow cytometry of A549-ACE2 cells at 3 days post-transduction with 30 or 100 uL SARS-CoV-2 spike pseudotyped lentivirus. (**c**) Percent of EGFP+ cells at 3 days post-transduction with the indicated volume of virus and pseudotype in human liver Huh7.5-ACE2 cells, lung A549-ACE2 cells, and colon Caco-2 cells (*n* = 3 replicates, error bars are s.e.m.). (**d**) The maximum fold-change in viral transduction in each cell line of G614 Spike as compared to D614 Spike (error bars are s.e.m.).

After transduction with 4 different viral volumes, we waited 3 days and then performed flow cytometry to measure GFP expression (**Figure 2b**). We found in all 3 human cell lines at all viral doses that G614 S-Virus resulted in a greater number of transduced cells than D614 S-virus (**Figure 2c**). Lentivirus lacking an attachment protein resulted in negligible transduction (**Figure 2c**). With the G614 Spike variant, the maximum increase in viral transduction over the D614 variant was 2.4-fold for Caco-2 colon, 4.6-fold for A549-ACE2 lung, and 7.7-fold for Huh7.5-ACE2 liver (**Figure 2d**). To control for any potential differences in viral titer, we also measured viral RNA content by qPCR. We observed only a small difference between D614 and G614 pseudotyped viruses using 2 independent primer sets (average of 7% higher viral titer for D614), which may result in a slight underestimation of the increase in transduction efficacy of the G614 pseudotyped virus (**Supplementary Fig. 2**).

We next sought to understand the mechanism through which the G614 variant increases viral transduction of human cells. Like SARS-CoV-1, the SARS-CoV-2 Spike protein has both a receptor-binding domain and also a hydrophobic fusion polypeptide that is used after binding the receptor (e.g. ACE2) to fuse the viral and host cell membranes (Heald-Sargent and Gallagher, 2012) (**Figure 3a**). Initially we hypothesized that the increased viral transduction of the G614 variant may be due to enhanced binding of the ACE2 receptor. To determine if greater transduction efficiency results from increased affinity of Spike G614 to its receptor, we used bio-layer interferometry to measure the binding kinetics of the Spike protein with and without the variant. We observed similar binding profiles of soluble D614 and G614 Spike to immobilized hACE2 (**Figure 3b, c**). Binding was best represented by a 2:1 heterogeneous binding model, which reports two dissociation constants (K_D_): K_D1_ = 8.45nM and K_D2_ = 127nM for D614 variant, and K_D1_ = 18.0nM and K_D2_ = 92.7nM for G614 variant (**Table 1**). K_D1_ is consistent with previously published binding affinities of the Spike D614 -ACE2 interaction (Walls et al., 2020; Yi et al., 2020), and is similar for the Spike G614 variant. This suggests our observed transduction phenotype is independent of Spike protein affinity for the ACE2 receptor.

**Table 1.** SARS-CoV-2 Spike D614 and G614 variants have comparable binding affinities for hACE2. Values of kinetics assay from bio-layer interferometry with purified Spike and ACE2. A 2:1 model yields two K_D_ values as measured by K_dis_/K_on_ for each binding process.

|  | **K<sub>D</sub>1**<br>(nM) | **K<sub>on</sub>1**<br>(1/M * s) | **K<sub>dis</sub>1**<br>(1/s) | **K<sub>D</sub>2**<br>(nM) | **K<sub>on</sub>2**<br>(1/M * s) | **K<sub>dis</sub>2**<br>(1/s) |
| --- | --- | --- | --- | --- | --- | --- |
| **Spike D614** | 8.45 | 3.51x10<sup>4</sup> | 2.96x10<sup>-4</sup> | 127 | 1.59x10<sup>5</sup> | 2.01x10<sup>-2</sup> |
| **Spike G614** | 18.0 | 4.25x10<sup>4</sup> | 7.67x10<sup>-4</sup> | 92.7 | 2.22 x10<sup>5</sup> | 2.06x10<sup>-2</sup> |

**Figure 3.**
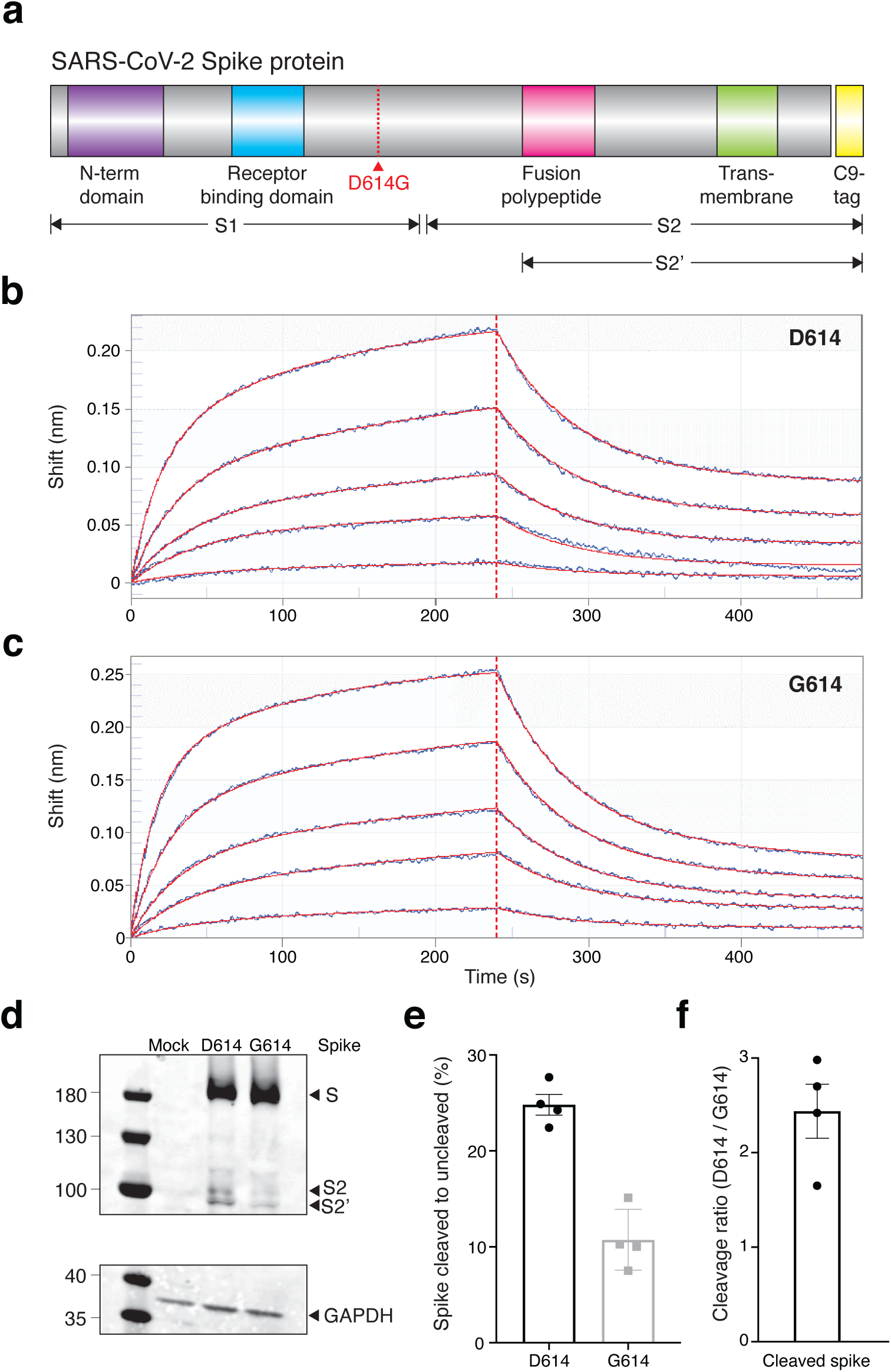
The SARS-CoV-2 Spike D614G variant displays similar ACE2 binding kinetics but altered proteolytic cleavage. (**a**) Schematic diagram of SARS-CoV-2 Spike protein structure with the added C9 affinity tag on the C-terminus. Spike cleavage fragments S1, S2, and S2’ are also indicated. (**b, c**) Association and dissociation binding curves of Spike D614 (**b**) and Spike G614 (**c**) with hACE2. Blue curves represent Spike protein binding profiles at 200 nM, 100 nM, 50 nM, 25 nM, and 6.25 nM. Red curves represent the best global fit using a 2:1 heterogeneous ligand model. (**d**) Western blot of total protein lysate from HEK293FT cells after transfection with D614 Spike, G614 Spike, or mock transfection. *(upper)* Detection of full-length Spike and cleavage fragments using an anti-C9 (rhodopsin) antibody. *(lower)* Detection of GAPDH via anti-GAPDH antibody. (**e**) Fraction of cleaved (S2 + S2’) to uncleaved (full-length) fragments for Spike D614 and G614 (*n* = 4 replicates, error bars are s.e.m.). (**f**) Fold-change in cleavage between Spike variants (D614 / G614) (*n* = 4 replicates, error bars are s.e.m.).

In order for SARS-CoV-2 to enter cells, the Spike protein must be cleaved at two sites by host proteases. It is thought that Spike must first be cleaved into S1 and S2 fragments, which exposes another cleavage site (Bestle et al., 2020; Hoffmann et al., 2020). The second cleavage event (creating the S2’ fragment) is thought to enable membrane fusion with the host cell. We transfected both D614 and G614 Spike variants into human HEK293FT cells to see if Spike cleavage might differ between these variants. Both constructs were tagged at their C-termini with a C9 tag to visualize full-length, S2, and S2’ fragments via western blot (**Figure 3d**). To measure cleavage, we quantified the ratio of cleaved Spike (S2 + S2’) to full-length Spike (**Figure 3e**). We found that the G614 variant is ∼2.5-fold more resistant to cleavage in the host cell than the D614 variant (**Figure 3f**). This suggests that the 2.4-to 7.7-fold increased transduction observed with G614 S-virus (**Figure 2d**) may be due to superior stability and resistance to cleavage of the G614 variant during Spike protein production and viral capsid assembly in host/producer cells.

Previous work showed that cleavage by the host protease furin at the Spike S1/S2 site in SARS-CoV-2 is essential for cell-cell fusion and viral entry (Hoffmann et al., 2020). To test for differences in furin-mediated cleavage, we performed *in vitro* digestion of both Spike variants after pull-down. We immunoprecipitated D614 and G614 Spike protein from HEK293FT cell lysates and then performed on-bead digestion using different concentrations of purified furin protease. Over a range of furin concentrations, we found that the G614 variant was more resistant to cleavage than the D614 variant (**Supplementary Fig. 3**). Importantly, the cleaved S2 and S2’ fragments might still be incorporated into new virions since they contain the required C-terminal transmembrane domain; however, they cannot functionally bind receptor due to lack of a N-terminal receptor binding domain (**Figure 3a**). Thus, the greater fraction of uncleaved G614 Spike may allow each newly-assembled virion to include more receptor binding-capable Spike protein.

Given the global efforts underway to develop a COVID-19 vaccine, we also sought to understand the impact of the Spike variant on immune responses. According to the World Health Organization, there are presently 126 COVID-19 vaccine candidates in preclinical development and 10 vaccine candidates in patient-enrolling clinical trials (https://www.who.int/publications/m/item/draft-landscape-of-covid-19-candidate-vaccines, accessed June 10th, 2020). Despite the tremendous diversity of vaccine formulations and delivery methods, many of the them utilize Spike sequences (RNA or DNA) or peptides and were developed prior to the emergence of the G614 variant. Using epitope prediction for common HLA alleles (Andreatta and Nielsen, 2016), we found that the G614 variant can alter predicted MHC binding (**Supplementary Fig. 4**). For example, the predicted binding for one high-affinity epitope decreased by nearly 4-fold (58nM for D614 versus 221nM for G614 with HLA-A*02:01). Although full-length Spike protein likely can produce many immunogenic peptides, several vaccines use only portions of Spike (Lurie et al., 2020; Yu et al., 2020) and thus it may be better to adapt vaccine efforts to the D614 variant given its global spread.

**Figure 4.**
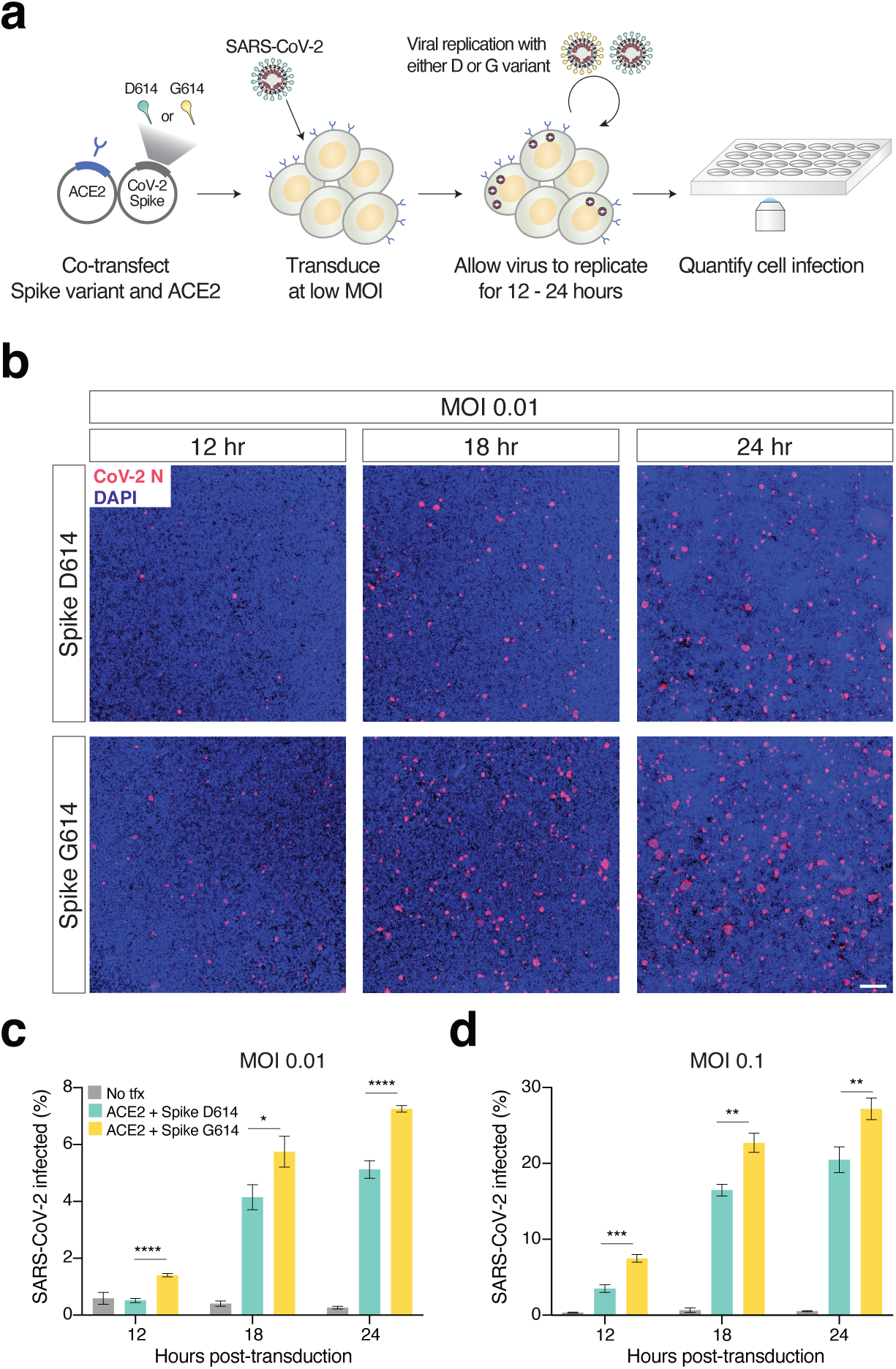
Increased infection of SARS-CoV-2 virus with the Spike D614G variant in human cells. (**a**) Schematic diagram of *trans*-complementation assay to assess the impact of SARS-CoV-2 Spike variants in an isogenic fashion. Plasmids containing a single Spike variant and ACE2 are co-transfected into HEK293T cells and then, after 24 hours, are infected with SARS-CoV-2 (Isolate USA-WA1/ 2020). (**b**) Representative images of HEK293T cells infected with SARS-CoV-2 (multiplicity of infection [MOI]: 0.01) and fixed at the indicated time point post-infection. The cells were stained with DAPI (blue) and an antibody for SARS-CoV-2 nucleocapsid protein (red). *Top row*: Transfection of ACE2 and SARS-CoV-2 Spike D614 plasmids. *Bottom row*: Transfection of ACE2 and SARS-CoV-2 Spike G614 plasmids. Scale bar: 1 mm. (**c, d**) Percent of SARS-CoV-2 infected cells at 12, 18, or 24 hours post-infection as measured by immunocytochemistry for SARS-CoV-2 nucleocapsid (N) protein and quantification using an imaging cytometer. A MOI of 0.01 in shown in panel **c** and a MOI of 0.1 is shown in panel **d**. * *p* ≤0.05, ** *p* ≤0.01, *** *p*≤0.001, **** *p* ≤ 0.0001.

To understand if the observed change in transduction efficiency that we found with our pseudotyped lentivirus also impacts full SARS-CoV-2 virus, we sought to develop an isogenic system for testing the Spike variant. Naturally-occurring isolates from patient samples that carry the Spike variant also carry a linked mutation in ORF1b, which makes it challenging to perform this experiment in an isogenic fashion. *In lieu* of a reverse genetics system to generate a SARS-CoV-2 variant and building on the observation by several groups that most cell lines require ACE2 overexpression for efficient SARS-CoV-2 infection (Hoffmann et al., 2020; Ou et al., 2020; Shang et al., 2020; Ziegler et al., 2020), we developed a novel *trans*-complementation assay in which we co-transfect either D614 or G614 Spike along with human ACE2 into HEK293T cells (**Figure 4a**). Twenty-four hours later, these cells were infected with SARS-CoV-2 at a low multiplicity of infection (MOI): In this manner, only transfected cells, which express ACE2 (and one of the Spike variants), can be readily infected by SARS-CoV-2.

We performed this experiment at two different MOIs (0.01 and 0.1) and measured differences in viral infection at 12, 18, and 24 hours post-infection (**Figure 4b, Supplementary Fig. 5**), as higher MOIs and longer infection periods may mask the contribution of transfected Spike in this assay. We found significant increases in infection with SARS-CoV-2 complemented with Spike G614 at all timepoints and all MOIs (**Figure 4c, d**). As a negative control, infected HEK293T cells without any prior transfection resulted in minimal SARS-CoV-2 infection (<1% in most cases). Using the *trans*-complementation assay, we have shown that introduction of just the Spike D614G point mutation increases infection using intact (replication-competent) SARS-CoV-2 virus.

In summary, we have demonstrated that the recent and now dominant mutation in the SARS-CoV-2 spike glycoprotein D614G increases the efficiency of cellular entry for the virus across a broad range of human cell types, including cells from lung, liver and colon. We demonstrated increased entry efficiency using both a pseudotyped lentiviral model system and also replication-competent SARS-CoV-2 virus. Given the concordance between the pseudotyped lentiviral system and SARS-CoV-2 virus, this suggests that changes in Spike protein are well represented using the pseudovirus, which should enable a much broader group of laboratories to use and study Spike variants.

We also found that G614 Spike is more resistant to proteolytic cleavage during production of the protein in host cells, suggesting that replicated virus produced in human cells may be more infectious due to a greater proportion of functional (uncleaved) Spike protein per virion. Using bio-layer interferometry with purified Spike and ACE2 proteins, we showed that there is no difference in binding kinetics with the ACE2 receptor resulting from the D614G mutation.

Since our initial preprint, several other groups have now confirmed that the D614G results in greater infection efficiency (Hu et al., 2020; Ozono et al., 2020; Yurkovetskiy et al., 2020; Zhang et al., 2020). Despite the emerging consensus the G614 results in faster viral spread (Korber et al., 2020), it is still uncertain whether this will have a clinical impact on COVID-19 disease progression. Two studies that have examined potential differences in clinical severity or hospitalization rates did not see a correlation with Spike mutation status (Korber et al., 2020; Wagner et al., 2020), although one study found a small but not significant enrichment of G614 mutations among intensive care unit (ICU) patients (Korber et al., 2020). Given its rapid rise in human isolates and enhanced transduction across a broad spectrum of human cell types, the G614 variant merits careful consideration by biomedical researchers working on candidate therapies, such as those to modulate cellular proteases, and on vaccines that deliver Spike D614 nucleic acids or peptides.

## Acknowledgements

We thank the entire Sanjana laboratory for support and advice. We are grateful to T. Maniatis, M. Legut, D. Ekiert, R. Redler, K. McGhee, C. Lu, and M. Prober for help with this work. Z.D. is supported by an American Heart Association postdoctoral fellowship (grant no. 20POST35220040). Postdoctoral fellowship support for T.X.J. is provided by the NIH (grant no. R01AI123155). G.B. is supported by PEW Biomedical Scholars (PEW-00033055), Searle Scholars Program (SSP-2018-2737) and NIH grant (grant no. R01AI147131). B.R.t. is supported by the Marc Haas Foundation, the National Institutes of Health, and DARPA’s PREPARE Program (HR0011-20-2-0040). N.E.S. is supported by New York University and New York Genome Center startup funds, National Institutes of Health (NIH)/National Human Genome Research Institute (grant nos. R00HG008171, DP2HG010099), NIH/National Cancer Institute (grant no. R01CA218668), Defense Advanced Research Projects Agency (grant no. D18AP00053), the Sidney Kimmel Foundation, the Melanoma Research Alliance, and the Brain and Behavior Foundation.

## Author contributions

Z.D. and N.E.S. conceived the project and designed the study. Z.D. performed transduction experiments with pseudotyped lentivirus and Spike protein biochemistry. T.X.J. and B.R.t. performed and analyzed the SARS-CoV-2 infections. J.I. and G.B. performed and analyzed bio-layer interferometry kinetic binding assays. X.G. analyzed SARS-CoV-2 genomes from patient isolates. All authors contributed to drafting and reviewing the manuscript, provided feedback and approved the manuscript in its final form.

## Methods

### SARS-CoV-2 genome analyses

For temporal tracking of D614G mutations in SARS-CoV-2 genomes, we used the Nextstrain analysis tool (https://nextstrain.org/ncov) with data obtained from GISAID (https://www.gisaid.org/)(Hadfield et al., 2018; Shu and McCauley, 2017). With the Nextstrain webtool, we visualized 3,866 genomes using the “clock” layout with sample coloring based on Spike 614 mutation status.

All complete SARS-CoV-2 genomes submitted before June 2nd 2020 were obtained from GISAID. We excluded genomes classified by GISAID as low coverage and downloaded the remaining 23,755 high-coverage genomes. To classify each genome as D614 or G614, we flanked the mutation site with 11-nt of surrounding sequence context on each side and identified genomes matching either mutation. For 1,652 genomes, we could not identify the mutation site and excluded these from further analysis. For the remaining 22,103 genomes, we were able to uniquely classify them as D614 or G614. Case-fatality rate data was downloaded on June 3rd 2020 from the Johns Hopkins Coronavirus Resource Center (https://coronavirus.jhu.edu/data/mortality). For accurate estimation of D614G prevalence, we only included countries with at least 9 genomes in GISAID.

### COVID-19 patient quantitative PCR

Threshold cycle data and statistical test results for Sheffield quantitative PCR data from COVID-19 patients is from Korber *et al*. (2020)(Korber et al., 2020). Threshold cycle data and statistical test results for University of Washington (UW) quantitative PCR data from COVID-19 patients is from Wagner *et al*. (2020) (https://github.com/blab/ncov-D614G)(Wagner et al., 2020). For the Sheffield study, the reported threshold cycle was the median in each group. For the UW study, the reported threshold cycle was the mean in each group. The reported *p*-values were computed by the respective study authors using the Wilcoxon Rank Sum test.

### Cell culture

A549 cells were obtained from ATCC, HEK293FT cells were obtained from Thermo Scientific, and Huh-7.5 and Caco-2 were a kind gift of B. tenOever (Mt. Sinai). All cells were cultured in D10 media: Dulbecco’s Modified Eagle Medium (Caisson Labs) supplemented with 10% Serum Plus II Medium Supplement (Sigma-Aldrich). Cells were regularly passaged and tested for presence of mycoplasma contamination (MycoAlert Plus Mycoplasma Detection Kit, Lonza).

### Spike plasmid cloning and lentiviral production

To express the D614 Spike, we used an existing CMV-driven SARS-CoV-2 plasmid (pcDNA3.1-SARS2-Spike, Addgene 145032)(Shang et al., 2020). To express the G614 Spike, we cloned pcDNA3.1-SARS2-SpikeD614G using the Q5 site-directed mutagenesis kit (NEB E0554S) and the following primers: 5’-CTGTACCAGGgCGTGAATTGCAC-3’ and 5’-CACGGCCACCTGGTTGCT-3’.

To make spike-pseudotyped lentivirus, we co-transfected a d2EGFP-containing transfer plasmid (Addgene 138152) with accessory plasmid psPAX2 (Addgene 12260) and the pseudotyping plasmid (or omitted the pseudotyping plasmid to produce no-pseudotype lentivirus). Briefly, for each virus, a T-225 flask of 80% confluent HEK293T cells (Thermo) was transfected in OptiMEM (Thermo) using 25 μg of the transfer plasmid, 20 μg psPAX2, 22 μg spike plasmid, and 175 μl of linear Polyethylenimine (1 mg/ml) (Polysciences). After 6 hours, media was changed to D10 media, DMEM (Caisson Labs) with 10% Serum Plus II Medium Supplement (Sigma-Aldrich), with 1 % bovine serum albumin (Sigma) added to improve virus stability. After 60 hours, viral supernatants were harvested and centrifuged at 3,000 rpm at 4 °C for 10 min to pellet cell debris and filtered using 45 μm PVDF filters (CellTreat). The supernatant was then ultracentrifuged for 2 hours at 100,000g (Sorvall Lynx 6000) and the pellet resuspended overnight at 4 °C in PBS with 1% BSA.

### Quantitative PCR (qPCR) of Spike pseudoviruses

Viral RNA was isolated from 100 mL of 100x-concentrated Spike D614 or G614 pseudotyped lentiviruses using 500 mL Trizol (Thermo 15596026) and following the Zymo Direct-zol RNA MicroPrep kit protocol. RNA was eluted with 15 mL RNase-free water. The RNA was then diluted 1:50 and 2 mL were used to perform a one-step qPCR protocol using Luna Universal One-step qPCR kit (NEB). Two primer sets were used: 5’-CGCTATGTGGATACGCTGC-3’ and 5’-GCGAAAGTCCCGGAAAGGAG-3’ that amplify WPRE, and 5’-CGTGCAGCTCGCCGACCAC-3’ and 5’-CTTGTACAGCTCGTCCATGCC-3’ that amplify EGFP. qPCR was performed following the Luna Universal One-step qPCR kit protocol on a ViiA 384-well qPCR machine.

### Spike pseudovirus transductions

We plated 50,000 cells per well of a 48-well plate. The cells were transduced the following morning using the indicated pseudotyped lentiviral amounts plus media supplemented with polybrene 8 μg/mL to a final volume of 150 μL per well. The media was changed 8 hours post-transduction. The cells were analyzed by flow cytometry 72 hours post-transduction.

### ACE2 lentiviral cloning and ACE2 stable cell line overexpression

To generate pLenti-ACE2-Hygro, we amplified human ACE2 (hACE2) from pcDNA3.1-ACE2 (Addgene 1786) and cloned it into a lentiviral transfer pLEX vector carrying the hygromycin resistance gene using Gibson Assembly Master Mix (NEB E2611L). A 2A epitope tag was added to hACE2 at the C-terminus. Huh7.5-ACE2 and A549-ACE2 cell lines were generated by lentiviral transduction of ACE2. The protocol for lentiviral production was the same as above except we used the common lentiviral pseudotype (VSV-g) using plasmid pMD2.G (Addgene 12259). Transduced cells were selected with hygromycin at 50 ug/mL for Huh7.5-ACE2 and 500 ug/mL for A549-ACE2 for 10 days before use.

### Flow cytometry of transduced human cells

Cells were harvested and washed with Dulbecco’s phosphate-buffered saline (Caisson Labs) twice. Cell acquisition and sorting was performed using a Sony SH800S cell sorter with a 100 μm sorting chip. We used the following gating strategy: 1) We excluded the cell debris based on the forward and reverse scatter; 2) Doublets were excluded. For all samples, we recorded at least 5000 cells that pass the gating criteria described above. Gates to determine GFP+ cells were set based on control GFP-cells, where the percent of GFP+ cells was set as <0.5% (background level). Flow cytometry analyses were performed using FloJo v10.

### ACE2 binding kinetics using bio-layer interferometry

The kinetics of the D614 and G614 Spike protein variants with hACE2 were analyzed using bio-layer interferometry on an Octet system (ForteBio, Octet RED96). Recombinant His-tagged and biotinylated human ACE2 protein (Sino Biological, Cat # 10108-H08H-B) was immobilized on a Streptavidin (SA) coated sensor. Loaded sensors were dipped into recombinant SARS-Cov-2 His-tagged Spike protein (D614 or D614G, Sino Biological, Cat # 40591-V08H and 40591-V08H3).

All proteins were diluted in kinetics buffer (0.1% w/v BSA, 0.02% Tween-20 in 1x PBS). Sensors were equilibrated in kinetics buffer for ten minutes at room temperature preceding data acquisition, and experiments were performed at 30 °C. Prior to ligand load, a baseline level was established for 60 s. hACE2 was loaded onto the sensor at 2.5μg/mL for 180 s, followed by a sensor wash (180 s) and a second baseline establishment (60 s) in kinetics buffer. Analyte in concentrations ranging from 200nM to 6.25nM were associated for 240 s and dissociated for 240 s. To determine K_D_ values for each variant, a reference sensor with loaded ligand but no analyte was subtracted from the data before fitting. Data was fit using a 2:1 heterogeneous ligand model from association and dissociation rates. The analysis was carried out with Octet ForteBio Analysis 9.0 software. Using a 2:1 model, two K_D_s were obtained. The higher affinity K_D_s are consistent with previously published values, and the second K_D_ values may represent a small amount of non-specific binding.

### Protein expression of ACE2 and spike variants in human cells

HEK293FT cells were transiently transfected with equal amounts of spike or ACE2 vectors using PEI. Cells were collected 18-24 hours post-transfection with TrypLE (Thermo), washed twice with PBS (Caisson Labs) and lysed with TNE buffer (10 mM Tris-HCl, pH 7.4, 150 mM NaCl, 1mM EDTA, 1% Nonidet P-40) supplemented with protease inhibitor cocktail (Bimake B14001) for 1 hour on a rotator at 4°C. Cells lysates were spun for 10 min at 10,000 g at 4°C, and protein concentration was determined using the BCA assay (Thermo 23227). Whole cell lysates (10 μg protein per sample) were denatured in Tris-Glycine SDS sample buffer (Thermo LC2676) and loaded on a Novex 4-12% Tris-Glycine gel (Thermo XP04122BOX). PageRuler pre-stained protein ladder (Thermo 26616) was used to determine the protein size. The gel was run in 1x Tris-Glycine-SDS buffer (IBI Scientific IBI01160) for about 120 min at 120V. Protein transfer was performed using nitrocellulose membrane (BioRad 1620112) using prechilled 1x Tris-Glycine transfer buffer (Fisher LC3675) with 20% methanol for 100 min at 100V. Membranes were blocked with 5% skim milk dissolved in PBST (1x PBS + 1% Tween 20) at room temperature for 1 hour. Primary antibody incubations were performed overnight at 4°C using the following antibodies: rabbit anti-GAPDH 14C10 (0.1 μg/mL, Cell Signaling 2118S), mouse anti-rhodopsin antibody clone 1D4 (1 μg/mL, Novus NBP1-47602) which recognizes the C9-tag added to the Spike proteins. Following the primary antibody, the blots were incubated with IRDye 680RD donkey anti-rabbit (0.2 μg/mL, LI-COR 926-68073) or with IRDye 800CW donkey anti-mouse (0.2 μg/mL, LI-COR 926-32212) for 1 hour at room temperature. The blots were imaged using Odyssey CLx (LI-COR). Band intensity quantification was performed by first converting Odyssey multichannel TIFFs into 16-bit grayscale image (Fiji) and the then selecting lanes and bands in ImageLab 6.1 (BioRad). In ImageLab, background subtraction was applied uniformly across all lanes on the same gel.

### On-bead Furin digestion of Spike protein

We transiently transfected 10-cm plates with 80% confluent HEK293FT with 10 μg of either spike D614 or G614 using PEI. About 24 hours later, cells were collected and lysed with 800 mL TNE buffer (10 mM Tris-HCl, pH 7.4, 150 mM NaCl, 1mM EDTA, 1% Nonidet P-40) supplemented with protease inhibitor cocktail (Bimake B14001) for 1 hour on a rotator at 4°C. Cells lysates were spun for 10 min at 10,000 g at 4°C. Spike was immunoprecipitated using 2 μg C9 antibodies (Novus NBP1-47602) per sample and incubated on a rotator at 4°C for at least 4 hours.

Recombinant Protein G Sepharose 4B beads (Thermo 101241) were washed twice with 1 mL TNE buffer and then were added to the immunoprecipitated cell lysate and incubated on a rotor at 4°C for 2 hours. Beads were then spun using a prechilled centrifuge at 4°C for 1 min at 2,000 rpm and washed 3x with 1 mL TNE. After the final spin, the beads were washed twice with 1 mL of furin reaction buffer (100 mM HEPES pH 7.5, 1 mM CaCl2, 1 mM b-Mercaptoethanol). Finally, the beads were resuspended in 150 μL and split equally in microcentrifuge tubes. The indicated amount of furin protease (NEB P8077) was added per reaction tube in a final volume of 20 μL. The reaction was incubated at 37°C for 1 hour and was occasionally mixed by gently tapping the tubes. Then the beads were denatured in Tris-Glycine SDS sample buffer (Thermo LC2676) and incubated at 95°C for 5 min. Samples were then loaded on a Novex 4-12% Tris-Glycine gel (Thermo XP04122BOX). Western blotting was performed as described above using mouse anti-rhodopsin antibody clone 1D4 (1 μg/mL, Novus NBP1-47602) which recognizes the C9-tag added to the Spike proteins. Following the primary antibody, the blots were incubated with IRDye 800CW donkey anti-mouse (0.2 μg/mL, LI-COR 926-32212) for 1 hour at room temperature. The blots were imaged using Odyssey CLx (LI-COR). Band intensity quantification was performed by first converting Odyssey multichannel TIFFs into 16-bit grayscale image (Fiji) and the then selecting lanes and bands in ImageLab 6.1 (BioRad). In ImageLab, background subtraction was applied uniformly across all lanes on the same gel.

### Epitope prediction using NetMHC

Since 9mer epitopes are most commonly presented by MHC receptors(Sarkizova et al., 2020), we constructed all possible 9mers surrounding the D/G 614 site in the Spike protein. We predicted binding affinities for 5 common HLA-A alleles and 7 common HLA-B alleles using the NetMHC 4.0 prediction webserver(Andreatta and Nielsen, 2016) (http://www.cbs.dtu.dk/services/NetMHC/). For each peptide, we computed the difference in predicted affinity between the D614 and G614 variant using R/RStudio and visualized them using the `pheatmap` R package.

### SARS-CoV-2 trans-complementation assay

SARS-related coronavirus 2 (SARS-CoV-2), Isolate USA-WA1/ 2020 (NR-52281) was deposited by the Center for Disease Control and Prevention and obtained through BEI Resources, NIAID, NIH. SARS-CoV-2 was propagated in Vero E6 cells in DMEM supplemented with 2% FBS, 4.5 g/L D-glucose, 4 mM L-glutamine, 10 mM Non-Essential Amino Acids, 1 mM Sodium Pyruvate and 10 mM HEPES.

For the trans-complementation assay, we co-transfected pcDNA3.1-SARS2-Spike or pcDNA3.1-SARS2-SpikeD614G (see above) with pcDNA3.1-ACE2 (Addgene 1786) at a 1:1 ratio (Spike : ACE2) using Lipofectamine 2000 (Thermo) as per the manufacturer’s protocol. At specified time points (12, 18 or 24 hr post-infection), cells were fixed using 5% formaldehyde and immunostained for nucleocapsid (N) protein (clone 1C7C7, Center for Therapeutic Antibody Discovery at the Icahn School of Medicine at Mount Sinai) with DAPI to stain nuclei. Full wells were imaged and quantified for SARS-CoV-2 infected cells using a Celigo imaging cytometer (Nexcelom Biosciences). All infections with SARS-CoV-2 were performed with 6 biological replicates.

### Statistical analysis

Data analysis was performed using R/Rstudio 3.6.1 and GraphPad Prism 8 (GraphPad Software Inc.). Specific statistical analysis methods are described in the figure legends where results are presented. Values were considered statistically significant for *p* values below 0.05.

### Reagent availability

All plasmids cloned for this study will be available on Addgene.

## References

Andreatta, M., and Nielsen, M. (2016). Gapped sequence alignment using artificial neural networks: application to the MHC class I system. Bioinforma. Oxf. Engl. 32, 511–517.

Becerra-Flores, M., and Cardozo, T. (2020). SARS-CoV-2 viral spike G614 mutation exhibits higher case fatality rate. Int. J. Clin. Pract. e13525.

Bestle, D., Heindl, M.R., Limburg, H., Van, T.V.L., Pilgram, O., Moulton, H., Stein, D.A., Hardes, K., Eickmann, M., Dolnik, O., et al. (2020). TMPRSS2 and furin are both essential for proteolytic activation and spread of SARS-CoV-2 in human airway epithelial cells and provide promising drug targets. BioRxiv 2020.04.15.042085.

Bhattacharyya, C., Das, C., Ghosh, A., Singh, A.K., Mukherjee, S., Majumder, P.P., Basu, A., and Biswas, N.K. (2020). Global Spread of SARS-CoV-2 Subtype with Spike Protein Mutation D614G is Shaped by Human Genomic Variations that Regulate Expression of TMPRSS2 and MX1 Genes. BioRxiv 2020.05.04.075911.

Dorp, L. van, Richard, D., Tan, C.C., Shaw, L.P., Acman, M., and Balloux, F. (2020). No evidence for increased transmissibility from recurrent mutations in SARS-CoV-2. BioRxiv 2020.05.21.108506.

Hadfield, J., Megill, C., Bell, S.M., Huddleston, J., Potter, B., Callender, C., Sagulenko, P., Bedford, T., and Neher, R.A. (2018). Nextstrain: real-time tracking of pathogen evolution. Bioinforma. Oxf. Engl. 34, 4121–4123.

Heald-Sargent, T., and Gallagher, T. (2012). Ready, set, fuse! The coronavirus spike protein and acquisition of fusion competence. Viruses 4, 557–580.

Hoffmann, M., Kleine-Weber, H., and Pöhlmann, S. (2020). A Multibasic Cleavage Site in the Spike Protein of SARS-CoV-2 Is Essential for Infection of Human Lung Cells. Mol. Cell 78, 779-784.e5.

Hu, J., He, C.-L., Gao, Q.-Z., Zhang, G.-J., Cao, X.-X., Long, Q.-X., Deng, H.-J., Huang, L.-Y., Chen, J., Wang, K., et al. (2020). The D614G mutation of SARS-CoV-2 spike protein enhances viral infectivity and decreases neutralization sensitivity to individual convalescent sera. BioRxiv 2020.06.20.161323.

Korber, B., Fischer, W.M., Gnanakaran, S., Yoon, H., Theiler, J., Abfalterer, W., Foley, B., Giorgi, E.E., Bhattacharya, T., Parker, M.D., et al. (2020). Spike mutation pipeline reveals the emergence of a more transmissible form of SARS-CoV-2. BioRxiv 2020.04.29.069054.

Li, W., Moore, M.J., Vasilieva, N., Sui, J., Wong, S.K., Berne, M.A., Somasundaran, M., Sullivan, J.L., Luzuriaga, K., Greenough, T.C., et al. (2003). Angiotensin-converting enzyme 2 is a functional receptor for the SARS coronavirus. Nature 426, 450–454.

Lurie, N., Saville, M., Hatchett, R., and Halton, J. (2020). Developing Covid-19 Vaccines at Pandemic Speed. N. Engl. J. Med. 382, 1969–1973.

Moore, M.J., Dorfman, T., Li, W., Wong, S.K., Li, Y., Kuhn, J.H., Coderre, J., Vasilieva, N., Han, Z., Greenough, T.C., et al. (2004). Retroviruses pseudotyped with the severe acute respiratory syndrome coronavirus spike protein efficiently infect cells expressing angiotensin-converting enzyme 2. J. Virol. 78, 10628–10635.

Ou, X., Liu, Y., Lei, X., Li, P., Mi, D., Ren, L., Guo, L., Guo, R., Chen, T., Hu, J., et al. (2020). Characterization of spike glycoprotein of SARS-CoV-2 on virus entry and its immune cross-reactivity with SARS-CoV. Nat. Commun. 11, 1620.

Ozono, S., Zhang, Y., Ode, H., Tan, T.S., Imai, K., Miyoshi, K., Kishigami, S., Ueno, T., Iwatani, Y., Suzuki, T., et al. (2020). Naturally mutated spike proteins of SARS-CoV-2 variants show differential levels of cell entry. BioRxiv 2020.06.15.151779.

Sarkizova, S., Klaeger, S., Le, P.M., Li, L.W., Oliveira, G., Keshishian, H., Hartigan, C.R., Zhang, W., Braun, D.A., Ligon, K.L., et al. (2020). A large peptidome dataset improves HLA class I epitope prediction across most of the human population. Nat. Biotechnol. 38, 199–209.

Shang, J., Ye, G., Shi, K., Wan, Y., Luo, C., Aihara, H., Geng, Q., Auerbach, A., and Li, F. (2020). Structural basis of receptor recognition by SARS-CoV-2. Nature 581, 221–224.

Shu, Y., and McCauley, J. (2017). GISAID: Global initiative on sharing all influenza data - from vision to reality. Euro Surveill. Bull. Eur. Sur Mal. Transm. Eur. Commun. Dis. Bull. 22.

Wagner, C., Roychoudhury, P., Hadfield, J., Hodcroft, E., Lee, J., Moncla, L., Müller, N., Behrens, C., Huang, M.- L., Mathias, P., et al. (2020). Comparing viral load and clinical outcomes in Washington State across D614G mutation in spike protein of SARS-CoV-2.

Walls, A.C., Park, Y.-J., Tortorici, M.A., Wall, A., McGuire, A.T., and Veesler, D. (2020). Structure, Function, and Antigenicity of the SARS-CoV-2 Spike Glycoprotein. Cell 181, 281-292.e6.

Yi, C., Sun, X., Ye, J., Ding, L., Liu, M., Yang, Z., Lu, X., Zhang, Y., Ma, L., Gu, W., et al. (2020). Key residues of the receptor binding motif in the spike protein of SARS-CoV-2 that interact with ACE2 and neutralizing antibodies. Cell. Mol. Immunol. 17, 621–630.

Yu, J., Tostanoski, L.H., Peter, L., Mercado, N.B., McMahan, K., Mahrokhian, S.H., Nkolola, J.P., Liu, J., Li, Z., Chandrashekar, A., et al. (2020). DNA vaccine protection against SARS-CoV-2 in rhesus macaques. Science.

Yurkovetskiy, L., Pascal, K.E., Tompkins-Tinch, C., Nyalile, T., Wang, Y., Baum, A., Diehl, W.E., Dauphin, A., Carbone, C., Veinotte, K., et al. (2020). SARS-CoV-2 Spike protein variant D614G increases infectivity and retains sensitivity to antibodies that target the receptor binding domain. BioRxiv 2020.07.04.187757.

Zhang, L., Jackson, C.B., Mou, H., Ojha, A., Rangarajan, E.S., Izard, T., Farzan, M., and Choe, H. (2020). The D614G mutation in the SARS-CoV-2 spike protein reduces S1 shedding and increases infectivity. BioRxiv 2020.06.12.148726.

Ziegler, C.G.K., Allon, S.J., Nyquist, S.K., Mbano, I.M., Miao, V.N., Tzouanas, C.N., Cao, Y., Yousif, A.S., Bals, J., Hauser, B.M., et al. (2020). SARS-CoV-2 Receptor ACE2 Is an Interferon-Stimulated Gene in Human Airway Epithelial Cells and Is Detected in Specific Cell Subsets across Tissues. Cell.

