## Supplementary Figures for "The Spike D614G mutation increases SARS-CoV-2 infection of multiple human cell types"

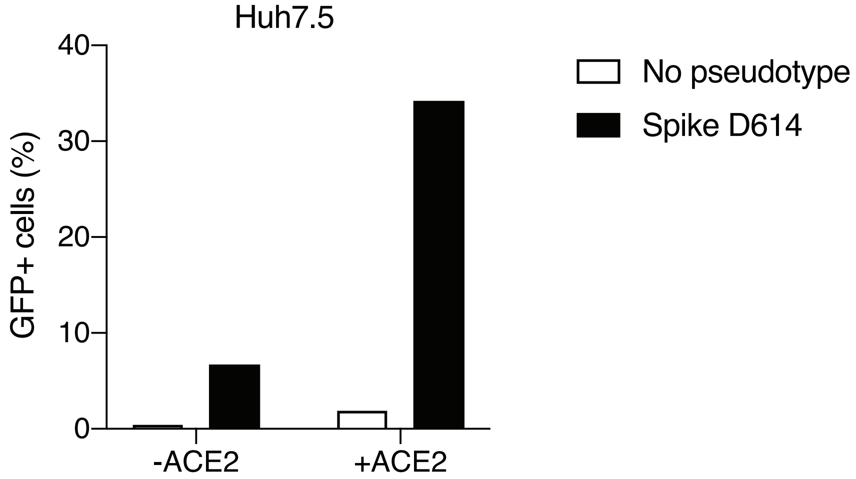


**Supplementary Figure 1. Increased transduction of SARS-CoV-2 Spike pseudotyped lentivirus in human cells that constitutively overexpress the human ACE2 receptor.** Percent of EGFP+ cells at 6 days post-transduction with 100 μL of supernatant SARS-CoV-2 spike (D614) pseudotyped lentivirus and unpseudotyped lentivirus in human liver Huh7.5 with and without ACE2 overexpression.

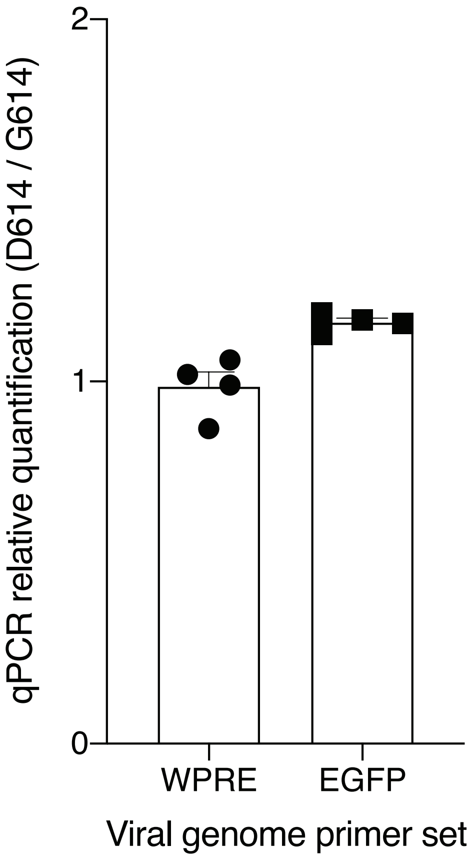


**Supplementary Figure 2. Quantitative PCR of viral RNA from SARS-CoV-2 Spike pseudotyped lentiviruses.** Relative quantification ($\Delta\Delta C_t$) of viral RNA from SARS-CoV-2 Spike pseudotyped lentiviruses. The primers amplify the Woodchuck Hepatitis Virus posttranscriptional regulatory element (WPRE) or the EGFP gene in the viral RNA genome.

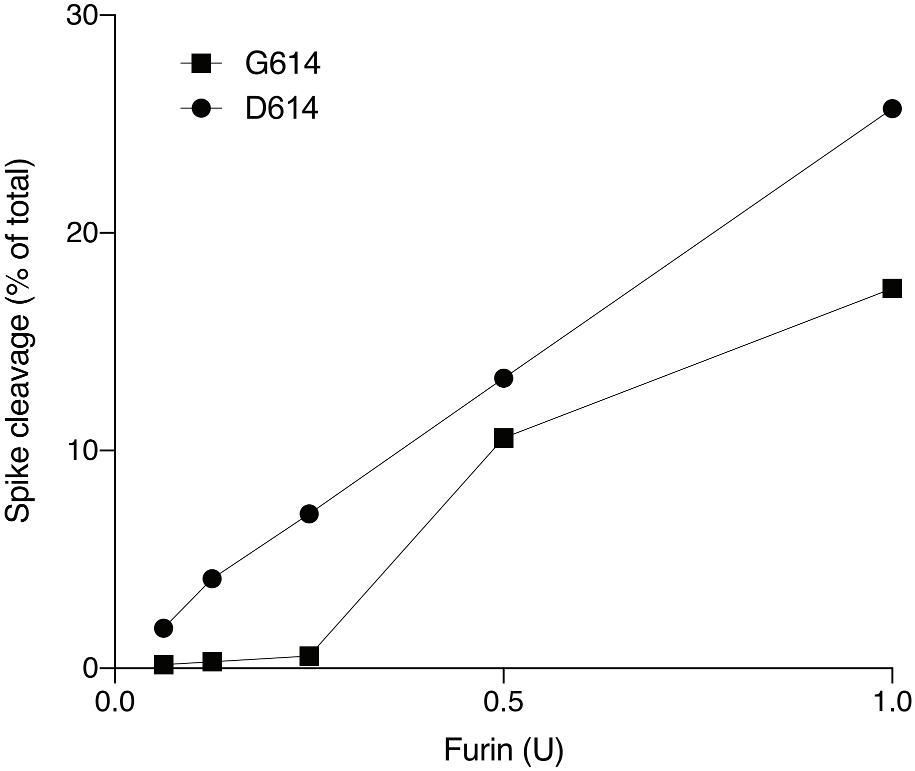


**Supplementary Figure 3. On-bead furin digestion of immunoprecipitated SARS-CoV-2 Spike variants.** Quantification of cleaved Spike (S2) fragment as a percent of total Spike after 1 hour digestion at 37°C with furin protease.

**a**

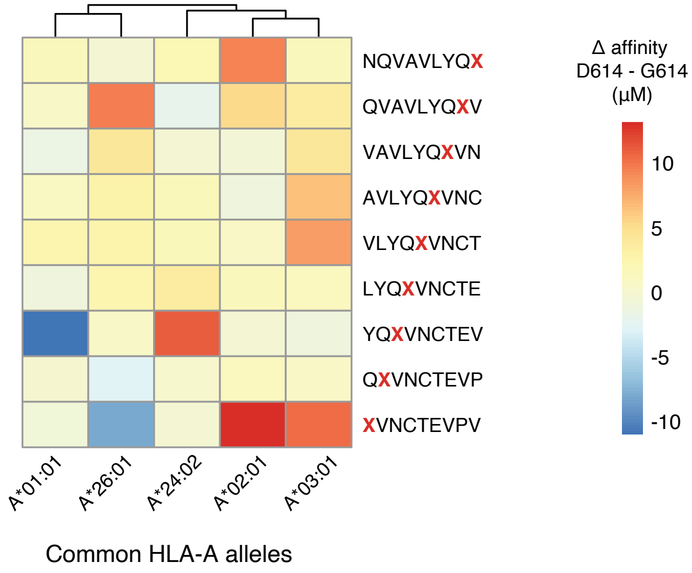


**b**

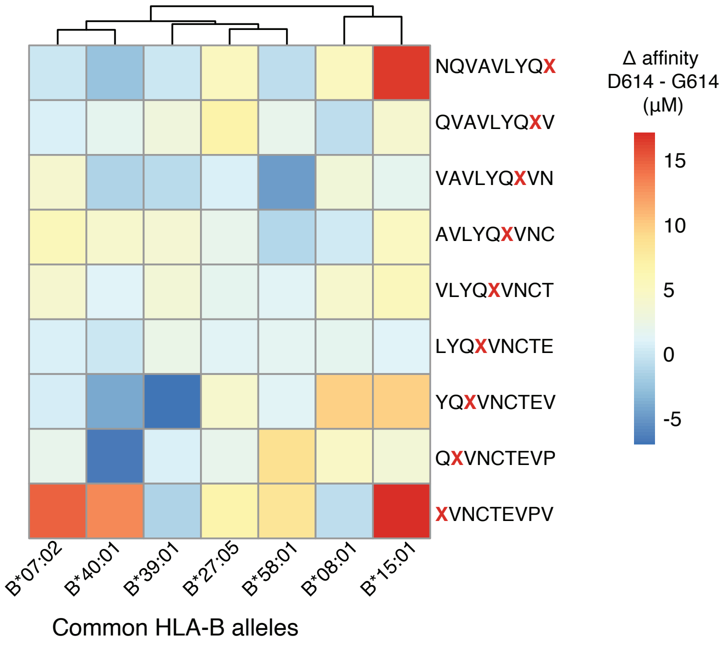


**Supplementary Figure 4. Change in MHC binding affinity for peptides near the D614G mutation in the SARS-CoV-2 spike protein.** Predicted change in binding affinity for common HLA-A alleles (a) and common HLA-B alleles (b). Predictions were computed using the NetMHC software package.

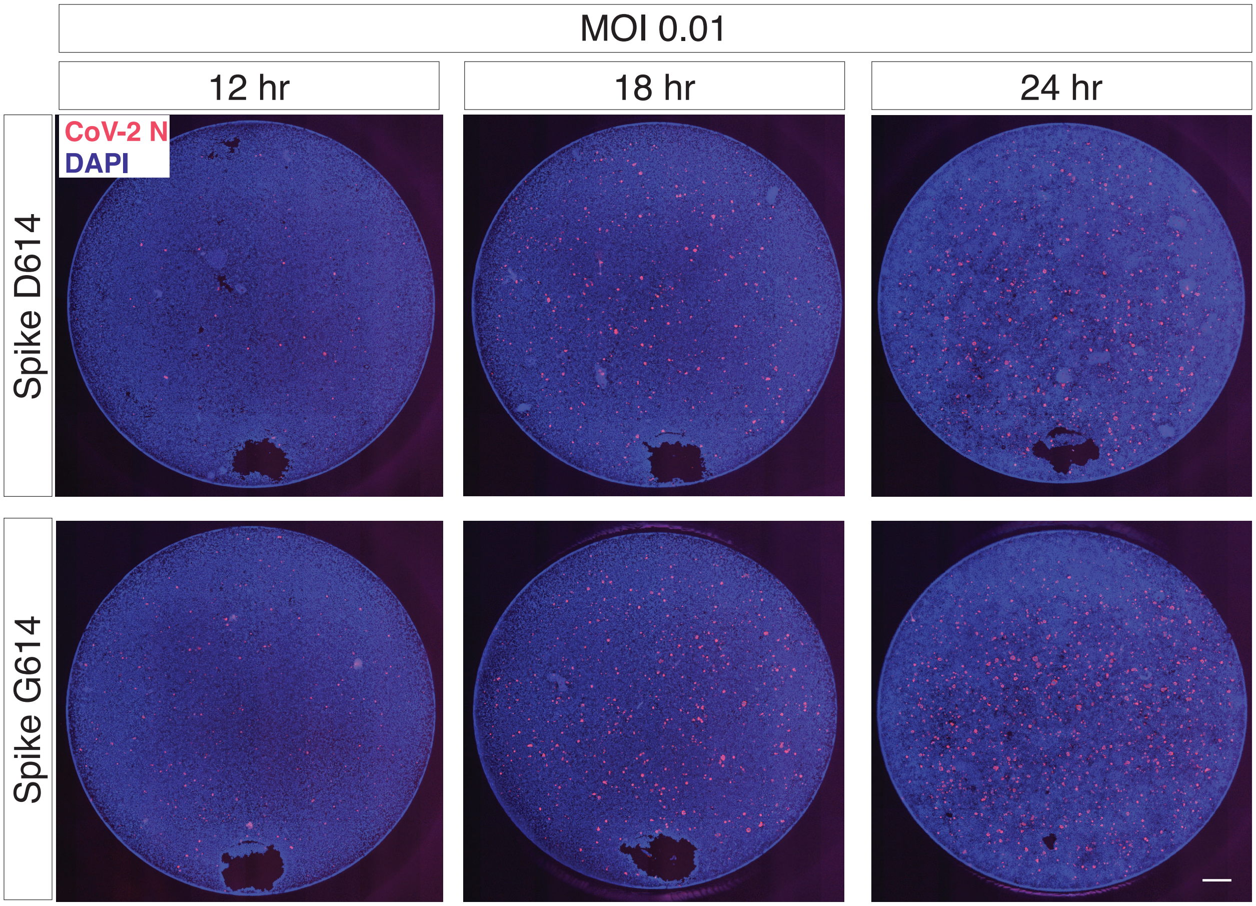


**Supplementary Figure 5. *Trans*-complementation of full SARS-CoV-2 virus with Spike G614 increases viral infection.** The regions shown in *Figure 4b* are representative sub-regions selected from the full images (entire well). The full images are shown here without any contrast adjustment for a single biological replicate ($n$ = 6 replicate infections for each time point-MOI combination). DAPI (nuclei) are shown in blue and SARS-CoV-2 nucleocapsid (N) protein is shown in red. Scale bar: 500 µm.
